# Excision of a 37-kb excision element as a circular plasmid in the cyanobacterium *Anabaena variabilis*

**DOI:** 10.64898/2025.12.09.693284

**Authors:** Brenda Pratte, Teresa Thiel

**Affiliations:** Department of Biology, University of Missouri-St. Louis, One University Dr., St. Louis, MO, 63130, USA

## Abstract

*Anabaena* (aka *Trichormus*) *variabilis* ATCC 29413 strain FD is a filamentous, heterocyst-forming cyanobacterium with a 6.36 Mb chromosome, three circular plasmids, A (366 kb), B (35.8 kb), C (301 kb), and a 37-kb excision element. This 37-kb element in *A. variabilis* ATCC 29413 strain FD was integrated into the *tRNA*^*cys*^ gene but was absent in the very closely related strains *A. variabilis* FSR and PNB. The 37-kb element was also excised as a circular molecule at a very low frequency compared to the integrated form. The 37-kb element has a partial copy of *tRNA*^*cys*^; therefore, integration produced a functional but genetically distinct copy of *tRNA*^*cys*^. Integration and excision are likely mediated by a putative integrase with a tyrosine recombinase domain encoded within the element, possibly using the duplicated copies of the 7-bp anticodon of the *tRNA*^*cys*^ as a recombination site. Like many other bacterial elements, the function of this element is unknown, but, like other well-characterized excision elements in heterocystous cyanobacteria, it does not appear to be important for survival.

## Introduction

*Anabaena (*aka *Trichormus) variabilis* ATCC 29413 is a filamentous, heterocyst-forming cyanobacterium. The original genome sequence of the *A. variabilis* ATCC 29413 variant strain FD (Currier and Wolk, 1979) consists of a 6.36 Mb chromosome, three circular plasmids A (366 kb), B (35.8 kb), and C (301 kb), and a 37-kb linear element (Thiel et al., 2014; GenBank assembly ID GCA_000204075). However, sequencing the original *A. variabilis* ATCC 29413 (not the FD variant),

Mardanov et al. identified a fourth plasmid, D (27 kb) (Mardanov et al., 2013), that was also seen in physical experiments (Lambert and Carr, 1982; Simon, 1978). Six cyanobacterial strains, originally cultured from the water fern *Azolla* (Zimmerman et al., 1989) that are nearly identical to *A. variabilis* ATCC 29413, also had plasmid D (Pratte and Thiel, 2021). In addition to the four circular plasmids, *A. variabilis* FD was reported to have a linear 37-kb element, whose mechanism of replication is unknown (Thiel, Pratte et al. 2014). The very similar strains *A. variabilis* FSR and *A. variabilis* PNB lack the 37-kb element (Pratte and Thiel, 2021). Thus, it appears that the 37-kb element is not essential.

Here, we further characterized the 37-kb linear element and determined that it integrates into a *tRNA* gene, which is a common site of integration in prokaryotes (Fouts, 2006; Reiter et al., 1989; Williams, 2002). We determined that in *A. variabilis* strain FD, it exists as a circular plasmid, although the integrated form, which predominates, may be required for replication.

## Results and Discussion

### Characterization of the 37-kb element

The genome sequence of the *A. variabilis* ATCC 29413 FD was reported to have a 37-kb linear element of unknown function and an unknown mechanism of replication (Thiel et al., 2014). Several other strains that we sequenced that are nearly identical to FD (*A. variabilis* ARAD, N2B, V5) have the same 37-kb element; however, the nearly identical strains *A. variabilis* PNB and *A. variabilis* FSR lack this element while retaining plasmids A–D (Thiel et al., 2014). When we sequenced these *Anabaena* genomes and resequenced the original *A. variabilis* ATCC 29413 FD strain, we found that at least some copies of the 37-kb linear element were in the chromosome between two *tRNA*^*cys*^ genes, one only a partial gene, that are flanked by genes *ava2569* and *psaB* (**Fig. 1A**). Similarly, in the finished genome of *A. variabilis* 0441 (Genbank CP047242) the 37-kb element is integrated in the chromosome (at nucleotides 4451261–4488411). Within this element in strain *A. variabilis* ATCC 29413 FD, there is a putative integrase gene, *ava-D0049*, which has a domain that is characteristic of tyrosine-type site-specific recombinases, such as bacteriophage lambda integrase (Nunes-Duby et al., 1998) and XisA, which removes the 11-kb excision element in the nitrogenase gene *nifD*, XisA, in *Nostoc* sp. PCC 7120 (Brusca et al., 1989). These site-specific recombinases target short *att* sites, often the anticodon loop in *tRNA* genes (Williams, 2002). The sequencing of thousands of bacterial genomes has revealed that *tRNA* genes are often the sites of insertion of genomic islands, which include, but are not limited to, pathogenicity islands (Dobrindt et al., 2004).

**Fig. 1.**
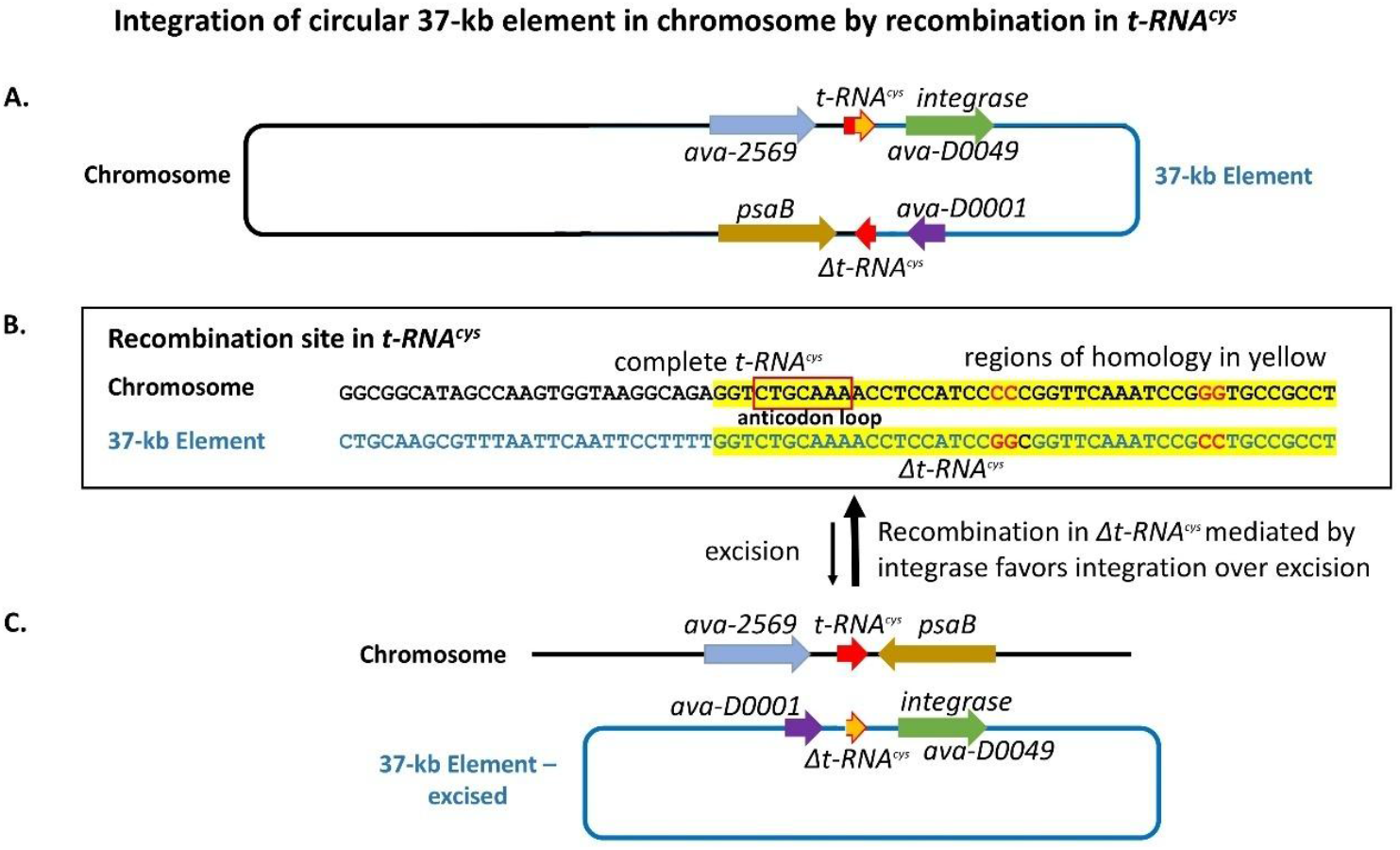
Model for the integration of the 37-kb element in the chromosome of *A. variabilis* ATCC 29413. A. The chromosome with the 37-kb element integrated in the *tRNA*^*cys*^ gene (not to scale). B. The region of homology in the *tRNA*^*cys*^ genes with the possible *att* recombination site in the anticodon loop in the red box. C. The chromosome and the circular form of the 37-kb element after excision.

We hypothesized that recombination between the conserved regions of the partial and complete copies of the *tRNA*^*cys*^ genes, mediated by integrase, allows integration and excision of the 37-kb element (**Fig. 1B**). Such an excision event should result in a 37-kb circular plasmid (**Fig. 1C**). We designed primers that would amplify the region of integration in the chromosome, with or without the 37-kb element, and the circular plasmid (**Fig. 2**). DNA from *A. variabilis* PNB, lacking the element, served as a control. PCR products were sequenced to verify the correct product, and the relative amount of each product was determined by qPCR. In *A. variabilis* ATCC 29413 FD, only about 0.5% of the chromosomes lacked the 37-kb element compared to 100% in *A. variabilis* PNB. About 93% of the total copies of the 37-kb element were in the chromosome in *A. variabilis* ATCC 29413 FD, while about 6% were in a circular form (**Table 1**).

**Fig. 2.**
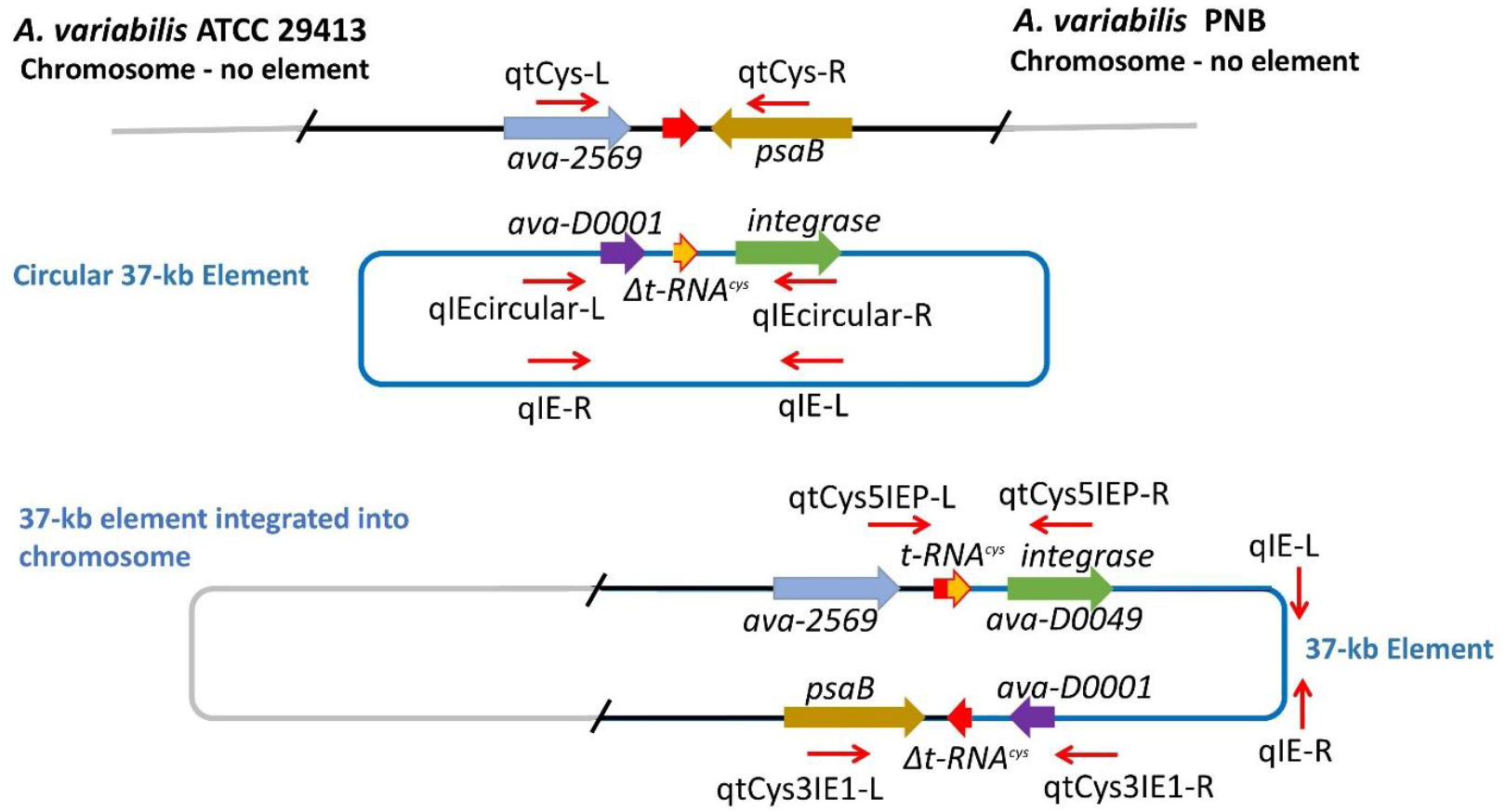
Primers and predicted products for integration or excision of the 37-kb element. The locations of the primers are shown by red arrows and their orientation indicates the predicted products based on integration or excision of the 37-kb element. The percentage of each DNA structure for *A. variabilis* ATCC 29413 or *A. variabilis* PNB is shown in Table 1.

**Table 1.** Distribution of element: integrated in the chromosome vs. the circular form.

|  | Chromosome<br>(no element)<sup>a</sup> | Element<sup>a</sup> |  |  |
| --- | --- | --- | --- | --- |
|  |  | Total | Chromosomal (integrated)/total | Circular/total |
| *A. variabilis* | 0.48% ± 0.07% | 100% | 92.8% ± 0.07% | 6.0% ± 1.1% |
| *Anabaena PNB* | 100% | 0 |  |  |

<sup>a</sup> Chromosomes lacking the 37-kb element were determined as the Cq value using qTCys-L/qTCys-R primers (see Fig. 2). The total amount of the 37-kb element was determined as the Cq value using primers qIE-L/qIE-R. The amount of circular 37-kb element was determined as the Cq value using primers qIEcircular-L/qIEcircular-R. The amount of the 37-kb element integrated in the chromosome was determined as the Cq value using primers qTCys3IE1-L/qTCys3IE1. Relative amounts of chromosomal (integrated) vs. circular forms of the 37-kb element were determined from $\Delta Cq$ values for the products made using the primers shown above. The $\Delta Cq$ values were converted to relative amounts ( $2^{\Delta Cq}$ ) and then converted to the appropriate percentage values as shown in the table.

**Table 2.**
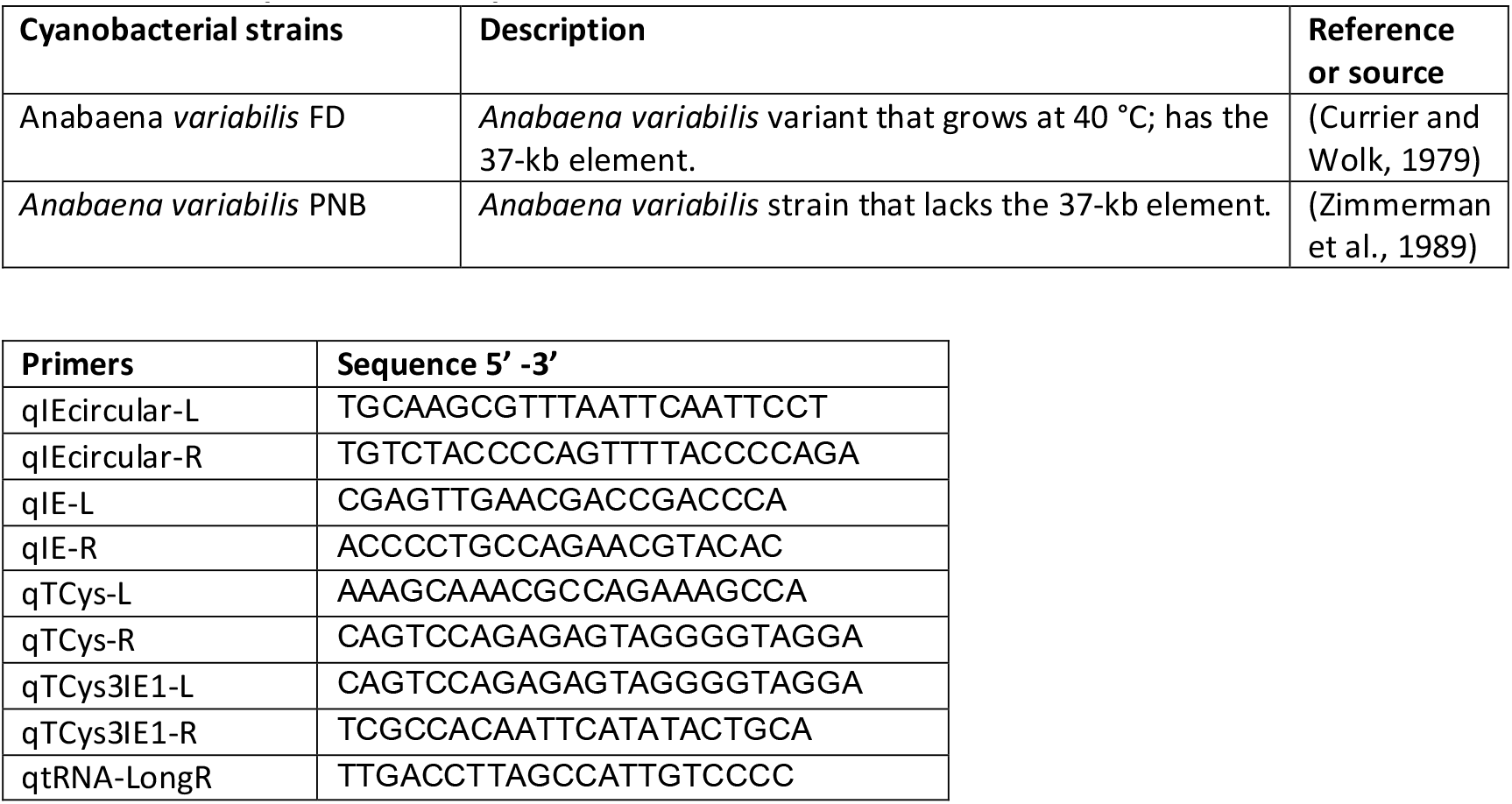
Strains, plasmids, and primers.

Thus, the integrated form of the 37-kb element, which is likely required for its replication, is the more common form in *A. variabilis* ATCC 29413 FD.

Since there is a single copy of *tRNA*^*cys*^, disruption of this gene by integration of the 37-kb plasmid would presumably be lethal. However, integration of the 37-kb plasmid into the chromosome creates a full copy of *tRNA*^*cys*^ consisting of the left part from the chromosome and the right part from the plasmid. Sequences of PCR products showed that this new gene differs from the native *tRNA*^*cys*^ gene in four nucleotides changing two Cs and two Gs (**Fig. 1B** and **Fig. 3**). This altered *tRNA*^*cys*^ is apparently functional, which is expected as the two pairs of mutations are compensatory and the predicted secondary structure is the same as that of the chromosomal gene, with a very similar ΔG (**Fig. 3**).

**Fig. 3.**
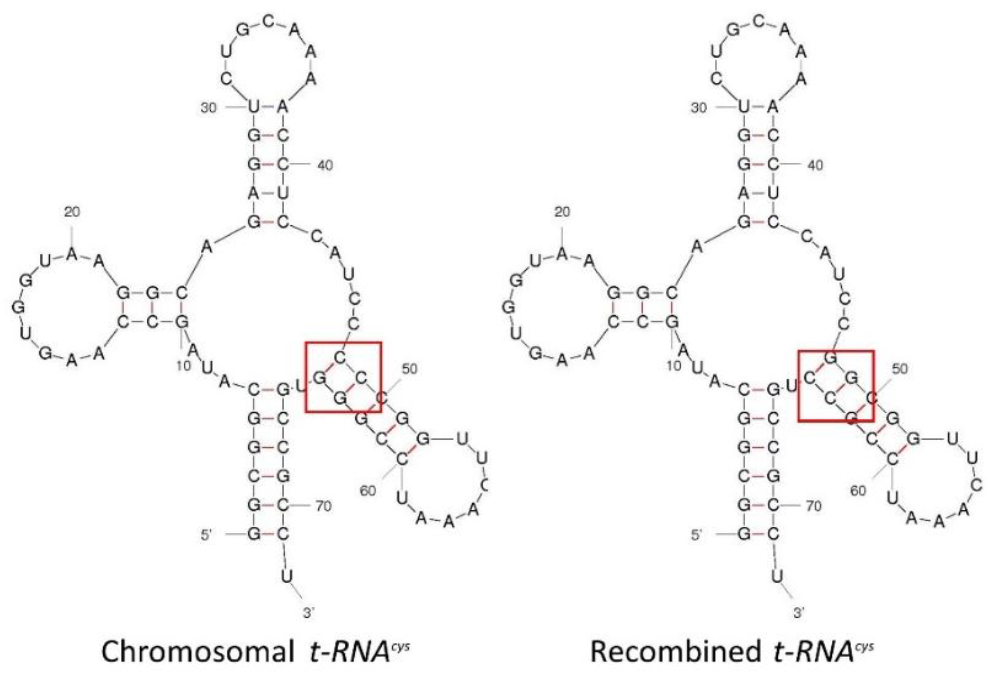
Predicted structures for *tRNA*^*cys*^. The chromosomal copy of *tRNA*^*cys*^ differs from the recombined *tRNA*^*cys*^ by four nucleotides, shown in the red boxes. However, the predicted structures are virtually identical, with a ΔG=−28.84 for the chromosomal *tRNA*^*cys*^ and ΔG= −29.34 for the recombined *tRNA*^*cys*^. A possible recombination site is the anticodon loop sequence from nucleotides 31–37 (5’-CUGCAAA-3’) (Williams, 2002).

Interestingly, the *A. variabilis* genome shows that *ava-2569*, upstream of *tRNA*^*cys*^, is itself part of a different 10,204-bp integrated element (from 3175621– 3185896 nucleotides in the genome), which duplicates the first 38 bp of *tRNA*^*cys*^. The 5’ region of *tRNA*^*cys*^ includes the putative *att* site in the anticodon loop (see Fig. 2B). Hence, the organization of the genome with the integrated 37-kb element is: 38-bp Δ*tRNA*^*cys*^—10-kb element—complete *tRNA*^*cys*^—37-kb element—45-bp Δ*tRNA*^*cys*^—*psaB*. Thus, there are two partial copies of *tRNA*^*cys*^ (38-bp N-terminal and 45-bp C-terminal) as well as the complete gene. Like the 37-kb element, the 10-kb element might be able to excise from the chromosome using its homology with the full *tRNA*^*cys*^, possibly at the anticodon loop site using a putative integrase (*ava-2575*). However, this integrase gene encodes a 147-aa protein, comprising only a tyrosine recombinase domain but lacking the N-terminal recombinase binding domain (identified using InterPro, (Blum et al., 2025)), which is very small for a functional integrase. The integrase in the 37-kb element is 368 aa, and the enzyme that excises the 11-kb excision element in the nitrogenase gene *nifD*, XisA (Brusca et al., 1989), is 472 aa. This suggests that the Ava-2575 integrase cannot excise the 10-kb element from the chromosome.

### Summary and conclusions

The 37-kb element that was first identified as linear extrachromosomal DNA in the JGI assembly of the genome of *A. variabilis* ATCC 29413 FD was labelled an excision element (Thiel et al., 2014). However, the element is integrated into the *tRNA*^*cys*^ gene of several very closely related strains of *A. variabilis*, except *A. variabilis* FSR and PNB, which lack the element (Pratte and Thiel, 2021). We also found the 37-kb element as an excised circular molecule in FD, but at a very low frequency compared to the integrated form. The 37-kb element has a partial copy of t*RNA*^*cys*^; therefore, integration produced a functional but genetically distinct copy of *tRNA*^*cys*^. The function, if any, of this newly identified excision element is unknown, but, like other excision elements in cyanobacteria, it does not appear to be important for survival.

## Methods

### Strains and growth conditions

*A. variabilis* 29413 strain FD and *A. variabilis* PNB were maintained on BG-11 agar medium (Rippka et al., 1979). Strains were grown photoautotrophically in liquid cultures in an eight-fold dilution of AA medium (AA/8)(Allen and Arnon, 1955) supplemented with 5 mM NH_4_Cl and 10 mM TES, pH 7.2, at 30 °C, with illumination at 50–80 μEinsteins m^−2^ s^−1^.

### DNA isolation/qPCR/sequencing for the 37-kb excision element

Genomic DNA was extracted from *A. variabilis* 29413 strain FD and *A. variabilis* PNB (a strain lacking the 37-kb excision element), grown in AA/8 (Allen and Arnon, 1955) with 5 mM NH_4_Cl and 10 mM TES pH 7.2, by vortexing cells with glass beads in the presence of phenol (Golden et al., 1985). Genomic DNA was purified with two phenol/chloroform/isoamyl alcohol extractions followed by a chloroform/isoamyl alcohol extraction before ethanol precipitation. qPCR reactions were performed using 4 ng of chromosomal DNA in a 10 µl reaction mixture using 5 pmol of gene-specific primers and 1x SsoAdvanced SYBR Green Supermix (BioRad). The following gene-specific primers were used: qIE-L/qIE-R, to amplify the incision element; qIEcircular-L/qIEcircular-R, to amplify the extrachromosomal circular IE; qTCys5IEP-L/qTCys5IEP-R, to amplify the 5’ end of *tRNA*^*cys*^ (next to the phage integrase side of the excision element); qTCys3IE1-L, qTCys3IE1-R, to amplify the 3’ end of *tRNA*^*cys*^ (next to the beginning sequence of the excision element); qTCys-L/qTCys-R, to amplify the genomic tCys (primers located outside the *tRNA*^*cys*^ gene) (see Fig. 2).

Sequencing of chromosomal DNA was performed to confirm the presence of the recombined form of *tRNA*^*cys*^. PCR products from the 5’ end and the 3’ end of the integrated incision element in the chromosome were amplified by PCR using primer sets qtCys5IEP-L/qIEcircular and qtRNA-LongR/qtCys3IE-R, respectively, confirming the presence of the recombined form of *tRNA*^*cys*^.

The 5’ and 3’ IE PCR products were purified using the QIAquick Gel Extraction Kit and sequenced using primers qIEcircular-R and qtRNA-LongR.

### Secondary structure of *tRNA*^***cys***^

Putative secondary structures for chromosomal and recombined forms of *tRNA*^*Cys*^ molecules, present with or without integration of the incision element in the chromosome, were determined using M-fold (Zuker, 2003).

## Acknowledgments

We are grateful to Jeff Elhai for his critique and helpful suggestions. This work was supported by the National Science Foundation grant MCB-1818298.

## Notes

### Competing Interest Statement

The authors have declared no competing interest.

## References

Allen, M.B., and Arnon, D.I. (1955). Studies on nitrogen-fixing blue-green algae. I. Growth and nitrogen fixation by Anabaena cylindrica Lemm. Plant Physiol 30, 366–372.

Blum, M., Andreeva, A., Florentino, L.C., Chuguransky, S.R., Grego, T., Hobbs, E., Pinto, B.L., Orr, A., Paysan-Lafosse, T., Ponamareva, I., et al. (2025). InterPro: the protein sequence classification resource in 2025. Nucleic Acids Res 53, D444–D456.

Brusca, J.S., Hale, M.A., Carrasco, C.D., and Golden, J.W. (1989). Excision of an 11-kilobase-pair DNA element from within the nifD gene in Anabaena variabilis heterocysts. J Bacteriol 171, 4138–4145.

Currier, T.C., and Wolk, C.P. (1979). Characteristics of Anabaena variabilis influencing plaque formation by cyanophage N-1. J Bacteriol 139, 88–92.

Dobrindt, U., Hochhut, B., Hentschel, U., and Hacker, J. (2004). Genomic islands in pathogenic and environmental microorganisms. Nat Rev Microbiol 2, 414–424.

Fouts, D.E. (2006). Phage_Finder: automated identification and classification of prophage regions in complete bacterial genome sequences. Nucleic Acids Res 34, 5839–5851.

Golden, J.W., Robinson, S.J., and Haselkorn, R. (1985). Rearrangement of nitrogen fixation genes during heterocyst differentiation in the cyanobacterium Anabaena. Nature 314, 419–423.

Lambert, G.R., and Carr, N.G. (1982). Rapid small-scale plasmid isolation by several methods from filamentous cyanobacteria. Arch Microbiol 33, 122–125.

Nunes-Duby, S.E., Kwon, H.J., Tirumalai, R.S., Ellenberger, T., and Landy, A. (1998). Similarities and differences among 105 members of the Int family of site-specific recombinases. Nucleic Acids Res 26, 391–406.

Pratte, B.S., and Thiel, T. (2021). Comparative genomic insights into culturable symbiotic cyanobacteria from the water fern Azolla. Microb Genom;7(6):000595. doi: 10.1099/mgen.0.000595. PMID: 34181515; PMCID: PMC8461463.

Reiter, W.-D., Palm, P., and Yeats, S. (1989). Transfer RNA genes frequently serve as integration sites for prokaryotic genetic elements. Nucleic Acids Research 17, 1907–1914.

Rippka, R., Deruelles, J., Waterbury, J.B., Herdman, M., and Stanier, R.Y. (1979). Generic assignments, strain histories and properties of pure cultures of cyanobacteria. J Gen Microbiol 111, 1–61.

Simon, R.D. (1978). Survey of extrachromosomal DNA found in the filamentous cyanobacteria. J Bacteriol 136: 414–418.

Thiel, T., Pratte, B.S., Zhong, J., Goodwin, L., Copeland, A., Lucas, S., Han, C., Pitluck, S., Land, M.L., Kyrpides, N.C., et al. (2014). Complete genome sequence of Anabaena variabilis ATCC 29413. Stand Genomic Sci. 2014 Jan 1;9(3):562–73. doi: 10.4056/sigs.3899418. PMID: 25197444; PMCID: PMC4148955.

Williams, K.P. (2002). Integration sites for genetic elements in prokaryotic tRNA and tmRNA genes: sublocation preference of integrase subfamilies. Nucleic acids research 30, 866–875.

Zimmerman, W.J.H., Rosen, B., and Lumpkin, T.A. (1989). Enzymatic, lectin, and morphological characterization and classification of presumptive cyanobionts from Azolla Lam. New Phytol 113, 497–503.

Zuker, M. (2003). Mfold web server for nucleic acid folding and hybridization prediction. Nucleic Acids Res 31, 3406–3415.

